# Whole-Brain Three-Dimensional Imaging of RNAs at Single-Cell Resolution

**DOI:** 10.1101/2022.12.28.521740

**Authors:** Shigeaki Kanatani, Judith C. Kreutzmann, Yue Li, Zoe West, Danai Vougesi Nikou, Jacob Lercke Skytte, Lea Lydolph Larsen, Daisuke H. Tanaka, Dagmara Kaczynska, Keishiro Fukumoto, Naofumi Uesaka, Tsutomu Tanabe, Ayako Miyakawa, Urmas Roostalu, Jacob Hecksher-Sørensen, Per Uhlén

## Abstract

Whole-brain three-dimensional (3D) imaging is desirable to obtain a comprehensive and unbiased view of architecture and neural circuitry. However, current spatial analytic methods for brain RNAs are limited to thin sections or small samples. Here, we combined multiple new techniques to develop TRIC-DISCO, a new pipeline that allows imaging of RNA spatial distributions in whole adult mouse brains. First, we developed Tris-mediated retention of *in situ* hybridization signal during clearing (TRIC), which produces highly transparent tissue while maintaining the RNA signal intensities. We then combined TRIC with DISCO clearing (TRIC-DISCO) by controlling temperature during the *in situ* hybridization chain reaction (isHCR) to ensure uniform whole-brain staining. This pipeline eliminates the requirements for both strict RNase-free environments and workflow-compatible RNase inhibitors. Our TRIC-DISCO pipeline enables simple and robust, single-cell, whole-brain, 3D imaging of transcriptional signatures, cell-identity markers, and noncoding RNAs across the entire brain.

## Introduction

Three-dimensional (3D) visualization of molecules on the whole-brain scale is essential to understand spatial relationships of cells and the circuits in which they participate (Ariel, 2017; Ueda et al., 2020). To date, multiple tissue clearing/staining techniques have been published, however, virtually all of them focus on the distribution of proteins rather than RNAs (Sylwestrak et al., 2016; Tian et al., 2021). Imaging RNA has several advantages over imaging proteins, including more flexible probe design and more rapid probe penetration into tissues (Sylwestrak *et al*., 2016). Although a few pioneering protocols have attempted to combine RNA imaging with tissue clearing (Sylwestrak *et al*., 2016; Tanaka et al., 2020), to our knowledge, none are applicable to larger tissues, such as entire mammalian brains.

*In situ* hybridization (ISH) is a common technique used to detect RNA. Traditionally, it uses digoxigenin-labeled RNA probes, in which signals are amplified by alkaline phosphatase using chromogens (Temsamani and Agrawal, 1996). However, enzyme-based signal amplification has been reported to generate a disproportionate brain signal gradient, higher near the brain surface than in the center, suggesting that it is not appropriate for 3D imaging (Sylwestrak *et al*., 2016). Recently, the *in situ* hybridization chain reaction (isHCR) has emerged as a new improvement of ISH. In isHCR, signal amplification is mediated by the HCR of fluorophore-labeled hairpin oligo DNA (Choi et al., 2014; Choi et al., 2018). We recently developed a technique (DIIFCO; diagnosing *in situ* immunofluorescence-labeled cleared oncosamples) that combines isHCR for RNA with organic solvent clearing (Tanaka *et al*., 2020). We have applied this technique in mouse embryos, brain sections, and clinical biopsies. Given the high demand for whole-brain RNA distribution analysis (Ueda *et al*., 2020), we also assessed the suitability of the DIIFCO pipeline for the whole adult mouse brain. However, due to low signal and/or low tissue transparency after clearing, we concluded that DIIFCO was unsuitable for visualizing RNA distribution across intact brains.

Here, we present a new technique, Tris buffer-mediated retention of *in situ* HCR signal during clearing (TRIC)-DISCO, which provides high tissue transparency after organic solvent (DISCO) clearing (Erturk et al., 2012; Renier et al., 2014) while not diminishing isHCR signals. We show that TRIC-DISCO is easy to perform and enables whole-brain imaging of a wide variety of transcripts at single-cell resolution, thereby providing a technique that can reveal 3D transcriptional landscapes deep inside intact brains.

## Results

### Reducing RNase contamination

Existing protocols for imaging RNAs in tissues are limited to ultra-thin sections or millimeter-thick samples because of issues with uneven probe penetration, low signal intensity, tissue transparency after tissue clearing, and poor reproducibility of imaging of thick samples. Therefore, we sought to overcome these shortcomings and allow the imaging of whole brains. First, we aimed to reduce the effect of RNase contamination, as this is a key technical issue in ISH, significantly affecting signal intensity and reproducibility, which is especially problematic for long and complicated protocols. We used the DIIFCO pipeline and removed all steps where RNase could be active until DNA hybridization was performed: low-percentage methanol treatment, blocking with donkey serum, and washing with aqueous buffers. Additionally, to protect RNA, we directly transferred the samples from 100% methanol to 50% formamide, which has been shown to inhibit RNase activity (Chomczynski, 1992). We next tested polyvinylsulfonic acid (PVSA), as it is a cost-effective RNase inhibitor, with an IC_50_ of 0.15 mg/mL against RNase A (Earl et al., 2018). Using a parvalbumin (*Pvalb*) probe in mouse cerebellum, we assessed three different PVSA concentrations: 3.8 mg/ml (∼85% RNase inhibition), 15 mg/ml (∼95% RNase inhibition), and 50 mg/ml (>99% RNase inhibition) in the amplification or hybridization buffer, or both (Fig. 1a and Supplementary Fig. 1). Because none of these significantly decreased *Pvalb* isHCR signal intensity (Fig. 1b; *n* = 3; one-way ANOVA, 3.8 mg/ml PVSA *F*_2,6_ = 1.586, *P* = 0.2799; 15 mg/ml PVSA *F*_2,6_ = 0.2738, *P* = 0.7695; 50 mg/ml PVSA *F*_2,6_ = 0.1584, *P* = 0.8569), we included 3.8 mg/ml (1:100 ratio) PVSA in our pipeline.

**Fig. 1.**
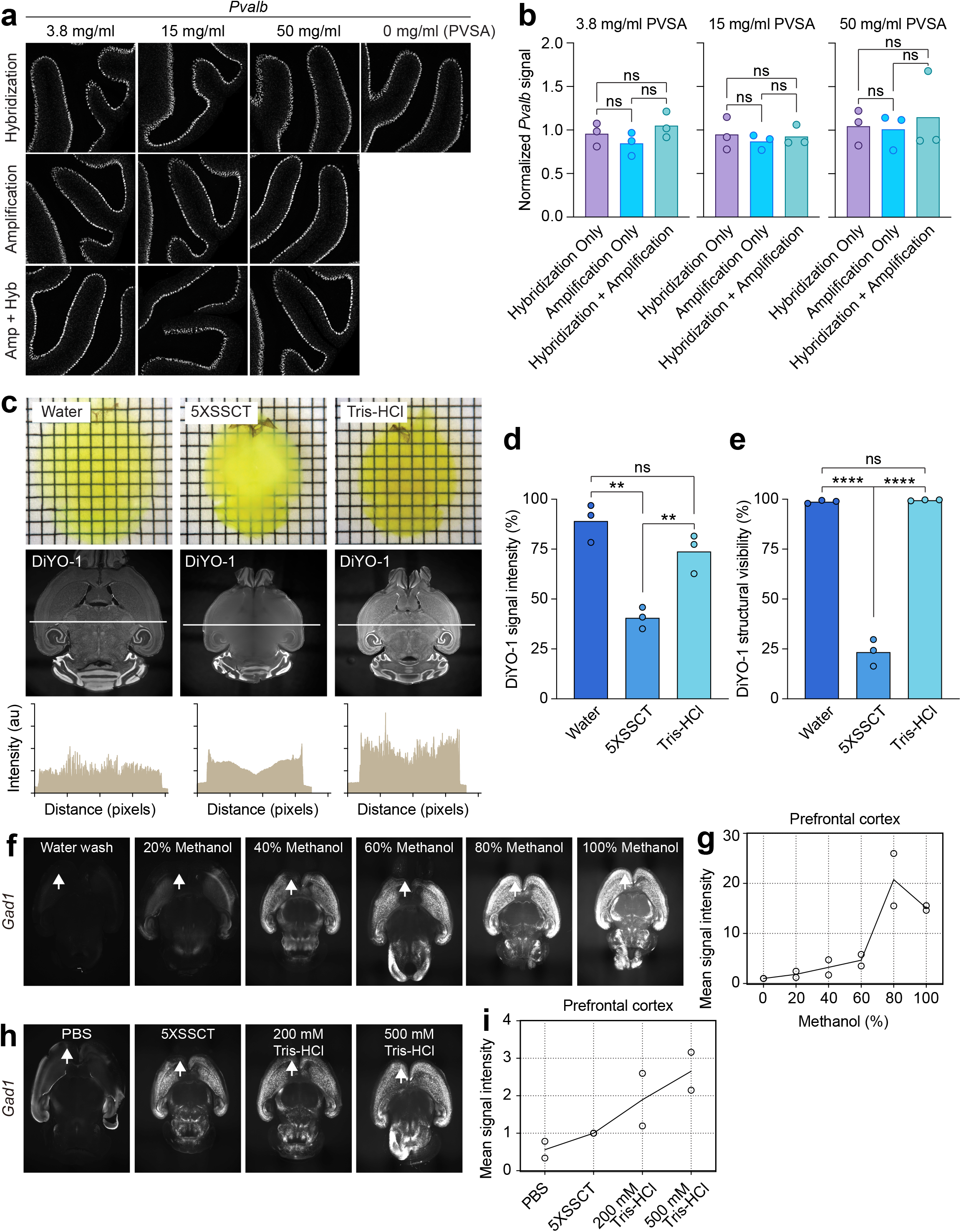
TRIC-DISCO bridges *in situ* HCR and DISCO tissue clearing. **a-b**, Representative images of isHCR for *Pvalb* in cerebellum sections using different concentrations of RNase inhibitor Polyvinyl sulfonic acid (PVSA) (3.8 mg/ml, 15 mg/ml, 50 mg/ml) in the amplification (Amp) or hybridization (Hyb) buffer, or both, (**a**) and the quantitative analysis of the signal intensity in Purkinje cells (**b**; *n* = 3, 3.8 mg/ml PVSA: Amp versus Hyb, *P* = 0.6242; Amp versus Amp+Hyb, *P* = 0.7077; Hyb versus Amp+Hyb, *P* = 0.2543; one-way ANOVA, *F*_2,6_ = 1.586, *P* = 0.2799; 15 mg/ml PVSA: Amp versus Hyb, *P* = 0.7621; Amp versus Amp+Hyb, *P* = 0.9769; Hyb versus Amp+Hyb, *P* = 0.8682; one-way ANOVA, *F*_2,6_ = 0.2738, *P* = 0.7695; 50 mg/ml PVSA: Amp versus Hyb, *P* = 0.9891; Amp versus Amp+Hyb, *P* = 0.9162; Hyb versus Amp+Hyb, *P* = 0.8540; one-way ANOVA, *F*_2,6_ = 0.1584, *P* = 0.8569). **c**, DISCO-cleared whole brains washed with water, 5XSSCT, or Tris-HCl and imaged on gridded paper (top; grid lines, 1 mm) or nuclear stained with DiYO-1 and imaged by light-sheet microscopy (middle) and quantification of signal intensity at the indicated line (bottom). au, arbitrary unit. **d-e**, Quantification of the DiYO-1 signal intensity (**d**; *n* = 3, water versus 5XSSCT, *P* = 0.0011; water versus Tris-HCl, *P* = 0.1544; 5XSSCT versus Tris-HCl, *P* = 0.0075; one-way ANOVA, *F*_2,6_ = 25.18, *P* = 0.0012) and the DiYO-1 structural visibility (**e**; *n* = 3, water versus 5XSSCT, *P* < 0.0001; water versus Tris-HCl, *P* = 0.9694; 5XSSCT versus Tris-HCl, *P* < 0.0001; one-way ANOVA, *F*_2,6_ = 377.4, *P* < 0.0001) in whole brains washed with water, 5XSSCT, or Tris-HCl. **f-i**, Representative images of isHCR for *Gad1* in whole brains (P0-1) washed for four days with water, 20%, 40%, 60%, 80%, or 100% methanol (**f**), or with PBS, 5XSSCT, 200 mM Tris-HCl, or 500 mM Tris-HCl (**h**), and the quantified signal intensity in the prefrontal cortex (**g, i**). The signal intensity was normalized to the mean intensity. All data are shown as the mean ± s.e.m. *n* denotes the number of biologically independent replicates. \*\**P* < 0.01, \*\*\*\**P* < 0.0001, ns, not significant by one-way ANOVA with Tukey’s post hoc comparison.

### Combining isHCR with organic solvent tissue clearing

Using DIIFCO, we occasionally observed that tissue transparency was insufficient for imaging the entire adult brain. Thus, we tested possible causes of insufficient clearing and found that prewashing the samples with pure water before clearing increased transparency (Fig. 1c). We further observed that the 5x saline-sodium citrate (5XSSC) buffer with Tween20 (5XSSCT) precipitated at high methanol concentrations, which resulted in insufficient clearing and significant loss of signal intensity (Fig. 1d; *n* = 3, Water versus 5XSSCT, *P* = 0.0011, one-way ANOVA, *F*_2,6_ = 25.18, *P* = 0.0012) and structure visibility (Fig. 1e; *n* = 3, Water versus 5XSSCT, *P* < 0.0001, one-way ANOVA, *F*_2,6_ = 377.4, *P* < 0.0001), assessed brain sections labeled with DiYO-1. The addition of water before clearing was futile, as doing so before organic solvent clearing removed virtually all isHCR signal and increased non-specific background (Fig. 1f). Thus, we searched for a formula to wash out salts prior to organic solvent tissue clearing while keeping the isHCR signal intact.

Using newborn (P0) mouse brains, we examined how dehydration during clearing reduced isHCR signal intensity. We found that washing with low methanol concentrations (0%, 20%, and 40%) for four days weakened the isHCR signal for Glutamate decarboxylase 1 (*Gad1*) and increased nonspecific background (Fig. 1f, arrows). In contrast, washing with higher methanol concentrations (60%, 80%, and 100%) for four days maintained the *Gad1* signal. Comparing signal intensities, we observed that the 60% methanol wash decreased the isHCR signal in the frontal cortex (Fig. 1g), whereas the 80% and 100% methanol washes gave stronger and more uniform signal intensities than the 60% methanol wash. These results suggest that a methanol concentration of at least 80% is needed to maintain isHCR signals during dehydration.

We next searched for buffers that could meet the requirement of solubility in >80% methanol. After testing many buffers, we found that tris(hydroxymethyl)aminomethane (Tris) was soluble in methanol. To test this modification, whole brains were incubated in Tris-HCl at pH 7.0. This treatment had a marked effect on the isHCR signal (Fig. 1f, arrows) while maintaining tissue transparency after clearing (Fig. 1c). Compared to 5XSSCT, Tris-HCl significantly enhanced signal intensity (Fig. 1d, *n* = 3, 5XSSCT versus Tris-HCl, *P* = 0.0075, one-way ANOVA, *F*_2,6_ = 25.18, *P* = 0.0012) and structural visibility (Fig. 1e, *n* = 3, 5XSSCT versus Tris-HCl, *P* < 0.0001, one-way ANOVA, *F*_2,6_ = 377.4, *P* < 0.0001). Importantly, while both 200 mM and 500 mM Tris-HCl buffer kept the overall isHCR signal intact (Fig. 1h, arrow), we observed reduced signal intensity in the frontal cortex with 200 mM Tris-HCl (Fig. 1i). Thus, we used 500 mM Tris-HCl buffer in subsequent experiments to wash out 5XSSCT before dehydration. We named this technique Tris-buffer-mediated retention of *in situ* HCR signal during clearing (TRIC). Notably, Tris-HCl buffer could not be used together with PVSA, as PVSA precipitated in methanol.

### Probe penetration and uniform staining

Although the penetration of DNA probes has been reported to be rapid (Sylwestrak *et al*., 2016), we obtained inconsistent results, particularly when a large number of DNA probes was used. While the surface of the brain showed intense staining, there was almost no signal in deeper regions, indicating that hairpin DNA was depleted on the surface. We hypothesized that applying a larger hairpin DNA volume would solve this problem, so we tested higher concentrations of hybridizing DNA probes and DNA hairpins. We compared the penetration of the *Gad1* probe when changing the total amount of hairpin DNA from 24 pmol to 96 pmol and found that this fourfold increase marginally improved penetration; however, this was not sufficient to cover the entire brain volume (Supplementary Fig. 2a,b). We also tested an eightfold increase of *Thy1* probe (192 pmol), but did not obtain sufficient signal intensity in deeper regions (Supplementary Fig. 2c,d, arrow). Thus, high concentrations of fluorescent hairpins cannot solve the issue of penetration, nor is doing so economically desirable.

Hence, we focused on the chemical reaction of the isHCR process to solve this issue. We assessed several approaches to achieving control over isHCR efficiency: changing buffer components, altering incubation time, adding pressure, and using centrifugation and electrophoreses, but without success. We also stained brain sections for *Pvalb* at multiple temperatures: 4°C, 20°C, 37°C, 55°C, and 70°C (Fig. 2a). This experiment revealed that 20°C and 37°C were optimal for isHCR (Fig. 2b); however, the *Pvalb* signal was still detected at 4°C. Intriguingly, when we assessed the effect of temperature for both *Thy1* (Fig. 2c) and the noncoding RNA *Malat1* (Fig. 2d), a dramatic decrease in surface signal was observed at 4°C. This effect likely occurred because the chemical process of isHCR at the tissue surface was heavily reduced at a low temperature, allowing a significant portion of hairpins to penetrate deeper into the tissue. Quantitative analysis showed a marked decrease in total signal intensity (Fig. 2e,f, *Thy1*: *n* = 2, 4°C versus 20°C, *P* = 0.0361, 4°C versus 37°C, *P* = 0.0207, one-way ANOVA, *F*_2,3_ = 19.05, *P* = 0.0197; *Malat1*: *n* = 3, 4°C versus 20°C, *P* < 0.0001, 4°C versus 37°C, *P* < 0.0001, one-way ANOVA, *F*_2,6_ = 279.7, *P* < 0.0001) and an increase in structural visibility (Fig. 2g,h, *Thy1*: *n* = 2, 4°C versus 20°C, *P* = 0.0022, 4°C versus 37°C, *P* = 0.0018, one-way ANOVA, *F*_2,3_ = 114.7, *P* = 0.0015; *Malat1*: *n* = 3, 4°C versus 20°C, *P* < 0.0001, 4°C versus 37°C, *P* < 0.0001, one-way ANOVA, *F*_2,6_ = 5774, *P* < 0.0001) at 4°C. From these results, we concluded that an increase in hairpin DNA at 4°C may allow uniform staining of transcripts across whole brains.

**Fig. 2.**
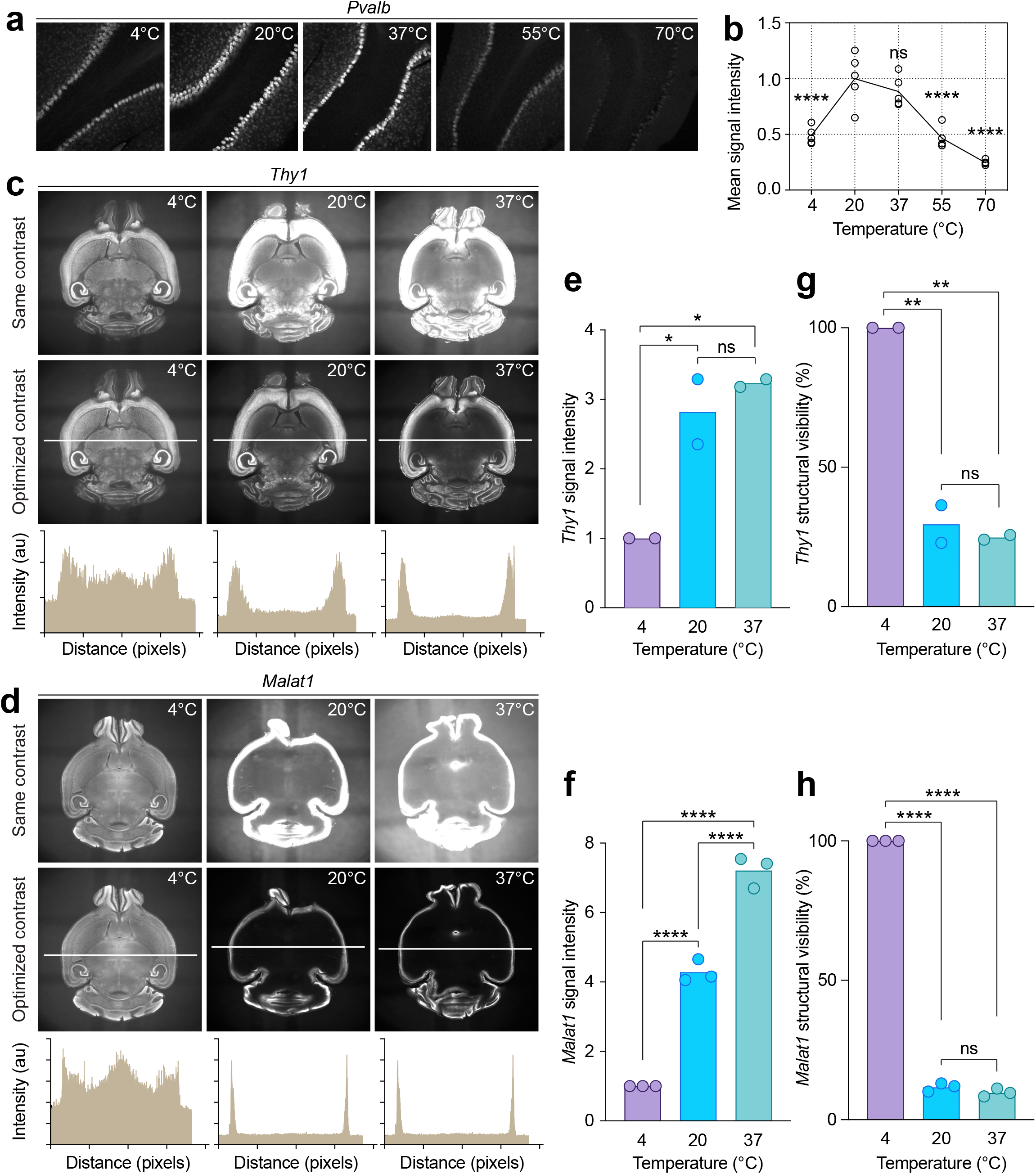
Temperature control of DNA probe penetration. **a-b**, Representative images of isHCR for *Pvalb* in cerebellum sections performed at temperatures: 4°C, 20°C, 37°C, 55°C, and 70°C (**a**) and the quantitative analysis of the signal intensity (**b**, *n* = 5, Control (Ctrl) was 20°C, Ctrl versus 4°C, *P* < 0.0001; Ctrl versus 37°C, *P* = 0.4584; Ctrl versus 55°C, *P* < 0.0001; Ctrl versus 70°C, *P* < 0.0001; one-way ANOVA, *F*_4,20_ = 28.23, *P* < 0.0001). The signal intensity was normalized to the mean intensity at 20°C. **c-f**, Representative images of isHCR for *Thy1* (**c**) and *Malat1* (**d**) in whole brains at 4°C, 20°C, or 37°C. The top panels show images normalized to 4°C. The middle panels show images normalized individually. The bottom plots show the quantification of the signal intensity at the indicated lines. au, arbitrary unit. **e**-**h**, Quantification of the signal intensity of isHCR for *Thy1* (**e**; *n* = 2, 4°C versus 20°C, *P* = 0.0361; 4°C versus 37°C, *P* = 0.0207; 20°C versus 37°C, *P* = 0.5913; one-way ANOVA, *F*_2,3_ = 19.05, *P* = 0.0197) and *Malat1* (**f**; *n* = 3, 4°C versus 20°C, *P* < 0.0001; 4°C versus 37°C, *P* < 0.0001; 20°C versus 37°C, *P* < 0.0001; one-way ANOVA, *F*_2,6_ = 279.7, *P* < 0.0001) and the structural visibility in whole brains isHCR for *Thy1* (**g**; *n* = 2, 4°C versus 20°C, *P* = 0.0022; 4°C versus 37°C, *P* = 0.0018; 20°C versus 37°C, *P* = 0.6955; one-way ANOVA, *F*_2,3_ = 114.7, *P* = 0.0015) and *Malat1* (**h**; *n* = 3, 4°C versus 20°C, *P* < 0.0001; 4°C versus 37°C, *P* < 0.0001; 20°C versus 37°C, *P* = 0.1812; one-way ANOVA, *F*_2,6_ = 5774, *P* < 0.0001). All data are shown as the mean ± s.e.m. *n* denotes the number of biologically independent replicates. \**P* < 0.05, \*\**P* < 0.01, \*\*\*\**P* < 0.0001, ns, not significant by one-way ANOVA with Dunnett’s (**b**) or Tukey’s (**e**-**h**) post hoc comparison.

### TRIC-DISCO enables uniform whole-brain RNA imaging at single-cell resolution

Finally, we performed 3D imaging of whole mouse brains using the entire TRIC-DISCO pipeline: PFA fixation, delipidation, bleaching, blocking, hybridization, wash, temperature-controlled hairpin probe incubation, TRIC processing and organic solvent clearing (Fig. 3a and Supplementary Protocol). We readily detected parvalubumin (*Pvalb*), somatostatin (*Sst*), Glutamate Decarboxylase 1 (*Gad1*), and nuclear staining with DiYO-1 (Fig. 3b and Supplementary Movies 1-2). We were able to recognize brain landmarks such as the neocortex and hippocampus; moreover, we observed overlapping and segregated expression patterns between *Pvalb, Sst, and Gad-1* across hippocampal neurons at single-cell resolution (Fig. 3c). TRIC-DISCO is also compatible with the green nuclear staining dye DiYO-1, which can be added during the DNA hybridization step. Taken together, our TRIC-DISCO pipeline provides a tool that allows integrative investigations of cellular structure and typology, noncoding RNA expression, and activity-regulated gene transcription in whole mouse brains.

**Fig. 3.**
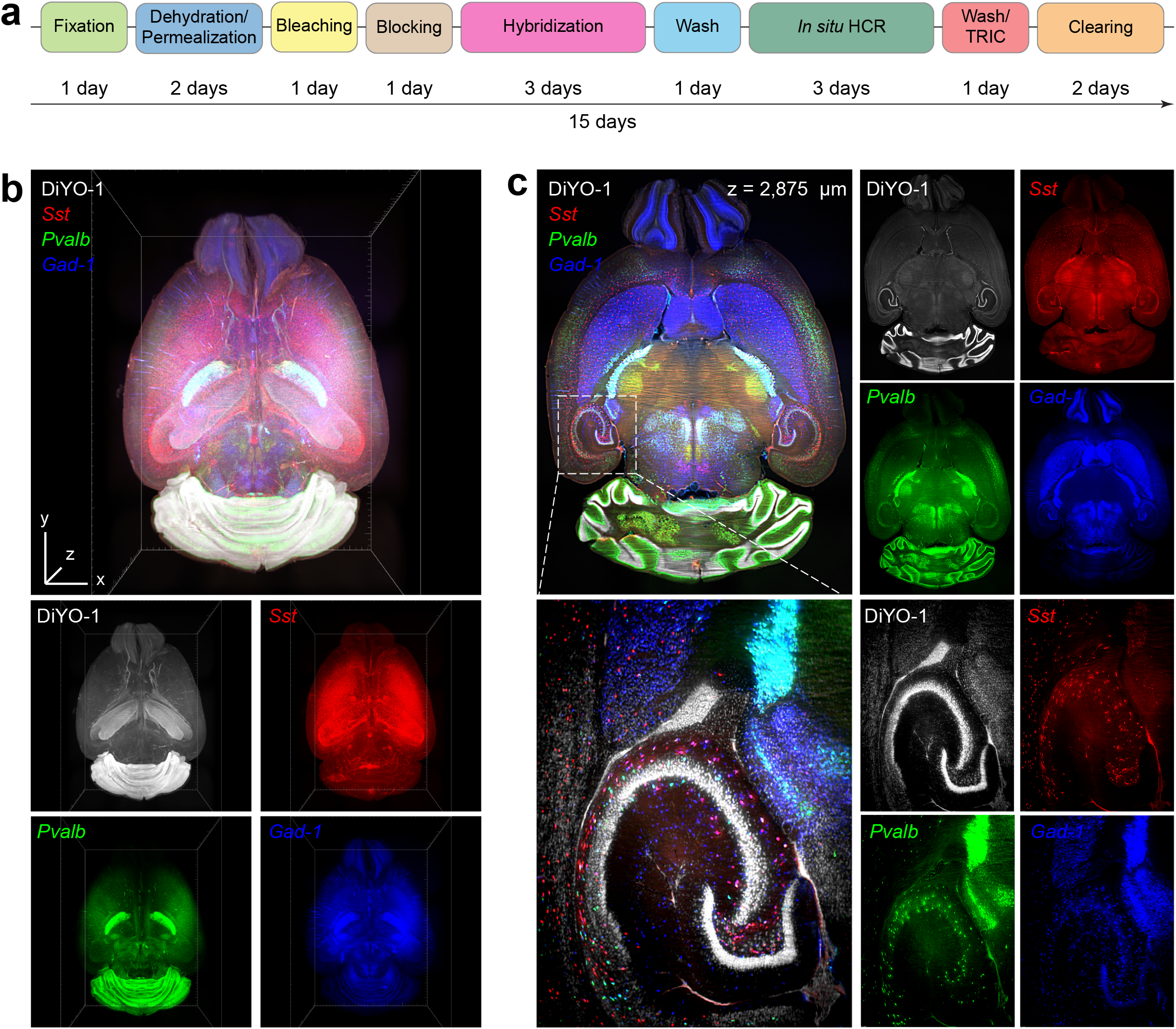
TRIC-DISCO pipeline enables analysis of RNA distributions in whole brains. **a**, Representative protocol for RNA detection in whole brains using TRIC-DISCO. The entire protocol requires 15 days. **b**, Representative images of isHCR for cortical interneuron transcripts: Somatostatin (*SST*, red), Parvalbumin (*Pvalb*, green), and Glutamate decarboxylase 1 (*Gad-1*, blue) in a whole brain from an eight-week-old adult mouse. Bounding box, 7,704 × 10,555 × 5,505 μm. **c**, Optical section at z = 2,875 μm of the same brain as in **b**. Magnified optical section of the hippocampus area at the indicated box.

### Application of TRIC-DISCO for a variety of genes

As our goal was to develop a simple, versatile, high-content, and scalable imaging method for whole brains, we sought to examine the expression patterns of multiple genes. We tested versatility using transcripts expressed in distinct cell types, including *Pvalb* and *Sst* for GABAergic inhibitory neurons, *Mfge8* and *Aldoc* for astrocytes, *Csflr* and *Hexb* for microglia, and *Sox10* and *Pdgfra* for oligodendrocytes and their precursors. Typical transcription patterns in the aforementioned cell types were readily detectable (Fig. 4a and Supplementary Movies 3-10). We also observed gene-specific enrichment in a variety of regions; for example, high cell density in the white matter for *Sox10* and *Pdgfra*, as well as dorsal enrichment of *Mfge8* (Zeisel et al., 2018).

**Fig. 4.**
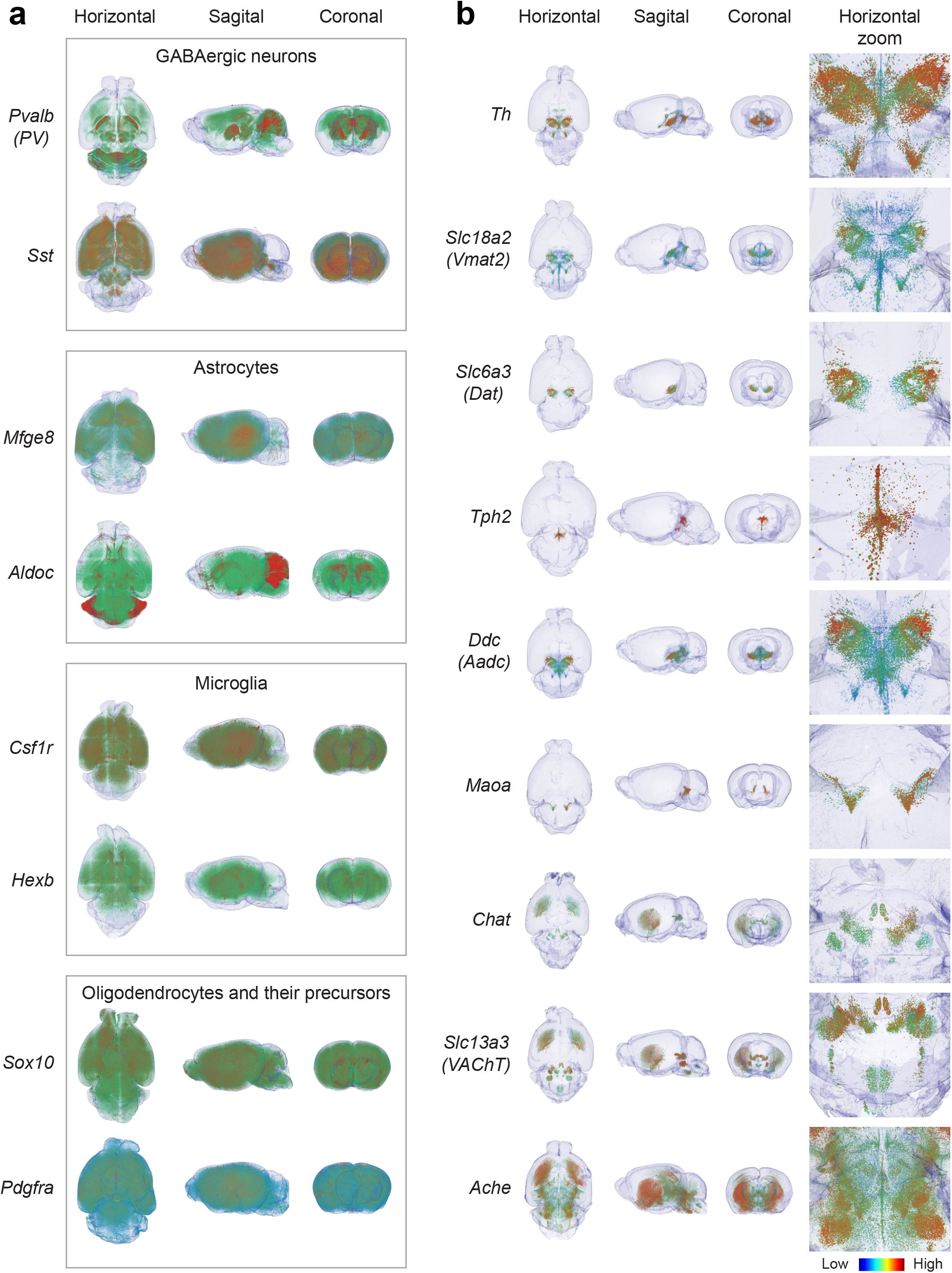
TRIC-DISCO can detect a variety of transcripts in whole brains. **a-b**, Horizontal, sagittal, and coronal views of isHCR for transcripts detected in GABAergic neurons (*Pvalb* (*PV*), *Sst*), astrocytes (*Mfge8, Aldoc*), microglia (*Csflr, Hexb*), oligodendrocytes and their precursors (*Sox10, Pdgfra*), (**a**), and neurotransmitters: *Th, Slc18a2* (*Vmat2*), *Slc6a3* (*Dat*), *Tph2, Ddc, Maoa, Chat, Slc13a3 (VAChT), Ache* (**b**) in whole brains from eight-week old mice. The horizontal view and zoom are from the ventral side.

We further analyzed the expression of genes related to neurotransmitters implicated in higher brain functions, such as dopamine, serotonin, and acetylcholine: *Th, Slc18a2, Slc6a3, Tph2, Ddc, Maoa, Chat, Slc13a3*, and *Ache*. TRIC-DISCO showed specific gene expression patterns in deep brain structures, such as basal ganglia and brain stem (Fig. 4b and Supplementary Movies 11-20). Volume rendering and magnification of 3D data revealed intricate transcriptional patterns of *Pvalb* and *Th* with single-cell resolution across the whole mouse brain (Supplementary Video 21-22). Taken together, these data document key steps toward the integrated investigation of cellular structure and typology, miRNA expression, and activity-regulated gene transcription within intact tissue volumes.

## Discussion

In this study, we improved the DIIFCO *in situ* hybridization method by solving multiple problems to make it applicable to whole adult mouse brains; we named it TRIC-DISCO. Its key improvements are: 1) temperature control of isHCR and probe penetration; 2) TRIC method that bridges isHCR with iDISCO clearing; and 3) an RNase-resistant and cost-effective workflow. By combining these techniques, TRIC-DISCO allows for the spatially uniform visualization of whole-brain transcripts at single-cell resolution. TRIC-DISCO is a simple, short, high-throughput protocol that can cover a wide range of molecules, allowing more reliable analyses of brain-wide phenomena.

Imaging in 3D using antibodies often shows a signal gradient from surface to deeper tissue layers (Renier *et al*., 2014). The uniform staining achieved with TRIC-DISCO allows the user to reduce or skip normalization image processing, which can be tedious, especially for large datasets such as whole brains. A further advantage of our method is the ability to design custom probes. Here, we tested a large number of custom DNA probes using the TRIC-DISCO pipeline (Supplementary Table 1) and confirmed their expression patterns (Perens and Hecksher-Sorensen, 2022). These include a noncoding RNA, as well as RNAs encoding proteins that are difficult to detect due to limited antibody availability. Our probe design method is simple, yet powerful enough to cover a large variety of RNAs. Further improvements in the field of probe design are likely to increase the signal intensity and success rate.

The TRIC-DISCO method is simple for beginners who have not performed isHCR due to its RNase-resistant workflow. Additionally, it is cost-efficient because it does not require special equipment and has a relatively low running cost, due to minimal DNA probes, fluorescent labeled hairpins, and commercially available reagents. Thus, TRIC-DISCO is available to many researchers in the field of neuroscience.

In this study, we showed that the efficiency of isHCR is temperature-dependent, with lower temperatures increasing the penetration of DNA hairpins. We further showed that low-temperature incubation during the isHCR process required significantly lower concentrations of DNA probes and hairpins for full penetration. Our approach provides unbiased whole-brain staining but may yield a somewhat weak signal from genes with lower expression levels. Therefore, there is room for technical improvements to handle full penetration and high signal intensity, for example, by adjusting the DNA probe length or by increasing the specificity of isHCR.

Time is a critical factor for high-throughput methods. Several steps in the TRIC-DISCO workflow can potentially be shortened to improve the protocol further. For example, we noticed that two days of sample incubation was sufficient for most transcripts used in this study; however, we used three days for some probes to achieve full isHCR amplification. Thus, our protocol describes three days of incubation to ensure that TRIC-DISCO works for all targets. The incubation times for different DNA probes will be further analyzed in future studies. Another step that can be shortened in the TRIC-DISCO protocol is the TRIC washing step, which was originally added to bridge isHCR and organic solvent clearing. Although this step could be removed by replacing 5XSSC with Tris-HCl buffer during isHCR, we prioritized the compatibility with isHCR as it is used worldwide. Together, these optimizations can reduce sample preparation time by as much as five days.

Techniques that loosen tissue barriers while keeping morphology intact are preferred for detecting different molecules deep inside tissues. It has previously been reported that urea-based buffers increase isHCR signal (Sinigaglia et al., 2018); this could potentially be applied by TRIC-DISCO to detect previously undetected RNAs. A combination of collagenase utilized in CUBIC-HistoVIsion (Susaki et al., 2020) or chemical treatment as utilized in SHANEL (Zhao et al., 2020) may contribute to the improvement of RNA detection in the future.

The development of new technologies is rapidly changing traditional low-throughput methods into comprehensive high-throughput approaches in neuroscience. Our TRIC-DISCO pipeline provides a new tool for performing spatial analyzes of RNAs in whole brains, which had not previously been feasible.

## Methods

### Mouse samples

Eight-week-old male C57BL/6JRj mice were purchased from Janvier Labs. P0-P1 mouse brains were collected from in-house bred C57BL/6JR mice. Mice were decapitated, and their brains were dissected after perfusion. Brains were immediately fixed in 4% paraformaldehyde in PBS overnight at 4°C and then directly transferred into 60%, 80%, 100%, 100% methanol (MeOH) and stored at - 20°C until use. All experiments were approved by Stockholm’s Ethical Committee North (ethical no. 16056-2017, 3644-2019).

### Probe design

Target RNA sequences were obtained from the NCBI gene database (https://www.ncbi.nlm.nih.gov/gene/). In cases of multiple variants, the longest sequence was selected and analyzed for sequence similarities using the Basic Local Alignment Search Tool (BLAST). Mouse genomic plus transcript (Mouse G+T) was used, and the option ‘Somewhat similar sequences (blastn)’ was selected. All other parameters were left as default. Similar sequences from the probe-target sequence, except for the sequence of the molecule and its variants, were excluded. Paired probes were designed in a tiling manner, with each pair having two base spaces. Each probe had the’ AA’ sequence’ between the split initiator and target sequence. The custom-written MATLAB code for designing probes is available on GitHub (https://github.com/uhlen-lab/TRIC-DISCO).

### Sample pretreatment (delipidation and bleaching)

Fixed samples were directly transferred to 100% dichloromethane (DCM) and incubated overnight. The samples were next washed twice with 100% MeOH and bleached in 5% H_2_O_2_ solution in MeOH overnight at 4°C. The samples were thereafter washed twice with 100% MeOH and stored at -20°C until further use.

### Sectioning for probe validation

Cryosections were prepared from samples that were stored in MeOH and directly transferred to 30% sucrose in PBS with PVSA (3.8 mg/ml) and incubated at 4°C until sinking. This step was repeated once. The samples were next embedded in O.C.T compound (Sakura fine Tek) and frozen at -80°C. Sections of 14 µm thickness were cut using a cryostat (CryoStar NX70, ThermoFisher) at the sagittal plane and stored at -20°C until further use.

Floating sections were prepared from samples that were stored in MeOH. The samples were washed with ethanol three times and cut with a vibratome (Leica VT1000ST, Leica, Germany) at 50 µm thickness and stored in 100% ethanol at -20°C until further use.

### isHCR on 2D sections

Custom-designed probes were initially tested on 2D sagittal cryosections or floating sections. The sections were washed twice in 30% formamide wash buffer for 10 min at RT. The sections were next blocked in 30% hybridization buffer for 30 min, at 37°C, in a moisture chamber, followed by hybridization in 30% hybridization buffer with 0.4 pmol/100 µl of probe (Supplementary Table 1) overnight at 37°C. The following day, the sections were washed with 30% formamide wash buffer three times at 37°C, followed by washing three times in 5XSSCT buffer at RT. For the isHCR reaction, the sections were incubated with the amplification buffer for 30 min at RT, followed by the amplification buffer with 6 pmol/100 µl DNA hairpin overnight at RT. The sections were thereafter washed three times in 5XSSCT buffer and mounted in 50% glycerol in 200 mM Tris-HCl buffer. Images were taken using a confocal microscope (Carl Zeiss LSM-980, Carl Zeiss LSM-900, or Olympus FV1000).

### Polyvinyl sulfonic acid (PVSA) concentration comparison

Brains stored in MeOH were washed twice in 100% ethanol (EtOH), and coronal sections (50 µm) were cut in 100% EtOH using a vibratome (Leica VT1000ST, Leica, Germany). The sections were stored in 100% EtOH until further use. When required, the brain sections were directly transferred to a hybridization wash buffer, and probe validation was performed. Different concentrations of PVSA (3.8 mg/ml, 15 mg/ml, 50 mg/ml) were added to the hybridization or HCR steps, and signal intensity was compared using probes for *Pvalb*.

### Whole brain isHCR

Delipidated and bleached samples that were stored in MeOH at -20°C were transferred to 50% formamide wash buffer and incubated at room temperature (RT) until they sank. The solution was changed twice. For blocking, the samples were transferred to 50% hybridization buffer and incubated overnight at 37°C with rotation. The samples were next transferred to 50% hybridization buffer with the respective DNA probe (1.6 pmol probe in 400 µl probe) and incubated for three days at 37°C with rotating. Samples were thereafter washed with 50% formamide wash buffer three times at 37°C, followed by three washes in 5XSSCT at RT. The samples were incubated in the amplification buffer overnight at RT or 4°C, followed by isHCR in the amplification buffer and fluorescently labeled DNA hairpins (24 pmol probe in 400 µl) for three days at RT or 4°C and washed three times in 5XSSCT buffer at RT, with the last wash overnight at RT, followed by the TRIC wash.

### Tris-mediated retention of signal intensity and clearing (TRIC)

Samples were washed in 500 mM Tris-HCl (pH 7.0) three times for one hour at RT to remove 5XSSCT and PVSA, with the last wash being performed overnight. The samples were next dehydrated in 100% MeOH three times for one hour at RT, with the last wash performed overnight. Following, dehydration and delipidation were performed by incubating the brains in a 66% DCM/33% MeOH solution for three hours, followed by 15 min of incubation in 100% DCM twice. Finally, the samples were transferred to dibenzyl ether (DBE) and incubated overnight. The DBE solution was changed once to increase transparency.

### Transparency test after isHCR

Samples were nuclear stained with DiYO-1 and washed with water, 5XSSCT, or Tris-HCl for four days. The solution was changed every day during the four-day incubation period. After washing, samples were dehydrated in a 60%, 80%, 100%, 100% MeOH series and delipidated and cleared, as described above. Transparency was quantified by two different criteria: distance until a uniform signal intensity could be detected (in % to total brain diameter) or cell structure visibility, with 100% indicating perfect visibility.

### Temperature test of isHCR

isHCR for *Pvalb* in sections was performed as described above. Floating sections were incubated in the amplification buffer at either 4°C, RT, 37°C, 55°C, or 70°C overnight. Whole brains were incubated at either 4°C, RT, or 37°C overnight. At 4°C, the samples were rotated at 20 rpm/min (Tube Revolver Rotator, ThermoFisher). The amplification volume was 400 µl for *Thy1* isHCR and 1,600 µl for *Malat1* isHCR.

### Dehydration condition test

Brains from P0-P1 animals were stained with a *Gad1* probe and washed three times with 200 mM Tris-HCl (pH 7.0). The samples were next incubated in one of the following conditions: pure water, 20%, 40%, 60%, 80%, 100% MeOH, 5XSSCT, 200 mM, 500 mM Tris-HCl, or PBS for four days at RT. The solution was changed every day during the four-day incubation period. After incubation, samples incubated in aqueous solutions or low concentrations of MeOH were dehydrated with 60%, 80%, 100%, and 100% MeOH solution for one hour. Samples incubated in 60% or 80% MeOH were dehydrated with a higher concentration of methanol (80% or 100%) for one hour, and then the 100% MeOH wash was repeated. Finally, all samples were treated with DCM and DBE, as mentioned in the TRIC-DISCO protocol. The intensity was normalized to the signal detected with water or 5XSSCT.

### Light sheet microscopy

Whole mouse brains were imaged with a LaVision Ultramicroscope II (MiltenyBiotec) or a Luxendo LCS-SPIM (Bruker) light-sheet microscope. When using the Luxendo LCS-SPIM light-sheet microscope, samples were imaged in quartz cuvettes filled with dibenzyl ether (DBE). A thin 2-3 mm layer of agarose, cleared according to the TRIC-DISCO protocol, was placed below the brain, in front of the olfactory bulbs, and behind the cerebellum to eliminate potential reflection from the cuvette surface. Samples were imaged at two wavelengths. Excitation at 561 nm and emission collected at 580-627 nm were used for tissue autofluorescence, and excitation at 642 nm and emission collected at 655-704 nm for the RNA-specific signal. Four-times magnification was used, and images were captured at 5 µm intervals, resulting in 1.46×1.46×5 µm voxel size. Single-sided illumination was used, with either the right or left lasers, depending on whether the right or left hemisphere was imaged. Exposure times were adjusted for individual samples. In post-processing, using the Luxendo-LCS SPIM software, individual image tiles were stitched and down-sampled to 5.85 µm X-Y resolution to enable faster data handling.

### Image processing

Image processing was performed using ImageJ (Rueden et al., 2017), Imaris (Bitplane), and Amira (Thermo Fisher Scientific Inc). The vessel signal was digitally cropped when unspecific vessel staining was observed outside the main expression region.

### Statistics

All quantitative data were expressed as the mean. The statistical test and definition *n* for each analysis are listed in the figure legends. Ordinary one-way ANOVA was used for comparisons across groups, and where significance was detected, a Dunnett’s or Tukey’s post hoc comparison was performed. No statistical methods were used to predetermine the sample size. Statistical tests were performed in GraphPad Prism V9 software (GraphPad Software, San Diego, CA, USA). Differences were considered significant at \**P* < 0.05, \*\**P* < 0.01, \*\*\**P* < 0.005, and \*\*\*\**P* < 0.001, and not significant (ns) at P ≥ 0.05. The experiments were not randomized, and investigators were not blinded to allocation during experiments and outcome assessment.

## Supporting information

Supplementary Protocol

## Acknowledgments

We would like to thank Maelle Bertho, Peter Löw, and Ole Kiehn from Karolinska Institutet, Stockholm, Sweden, for providing us with newborn mouse brains. This work was supported by the Swedish Research Council (grants 2017-00815 and 2021-03108 to P.U.), the Swedish Brain Foundation (grants FO2018-0209 and FO2020-0199 to P.U.), the Swedish Cancer Society (grants 19 0544 Pj, 19 0545 Us, and 22 2454 Pj to P.U.), and the Swedish Childhood Cancer Foundation (grants PR2020-0124 and PR2022-0111 to P.U.). The light-sheet microscopy infrastructure used in this research received grants from the Strategic Research Area in Neuroscience (StratNeuro) and the Strategic Research Area in Stem Cells and Regenerative Medicine (StratRegen), supported by the Swedish government.

## Notes

### Competing Interest Statement

The authors have declared no competing interest.

