## Supplementary Information for "Whole-Brain Three-Dimensional Imaging of RNAs at Single-Cell Resolution"

### TRIC-DISCO: Whole-Brain Imaging of RNA

#### Supplementary Figures

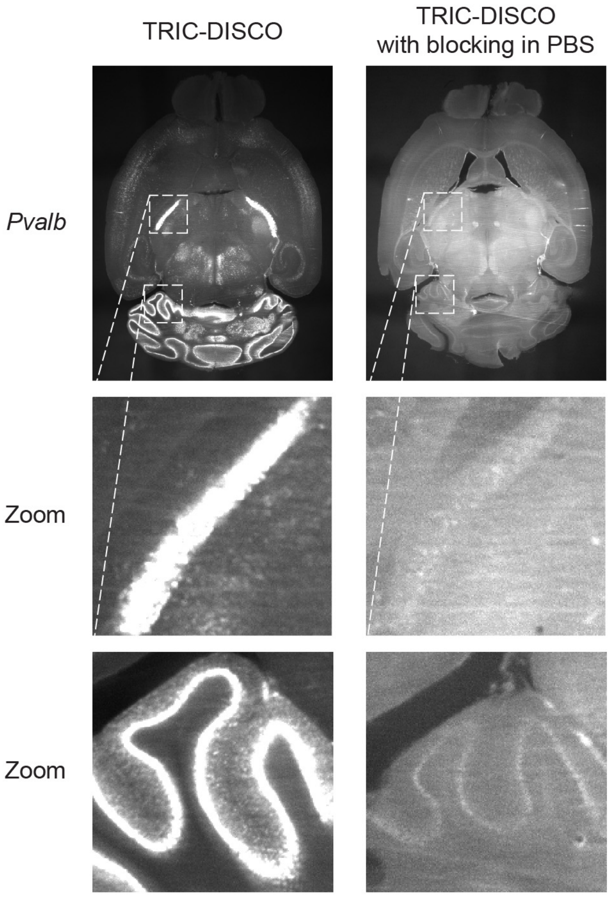

**Supplementary Figure 1 | Assessment of the signal intensity of isHCR using blocking.** Comparison of the signal intensity of isHCR for *Pvalb* using TRIC-DISCO or TRIC-DISCO with blocking solution (donkey serum in PBS without PVSA) for 30 min. Zoomed views of the indicated boxes.

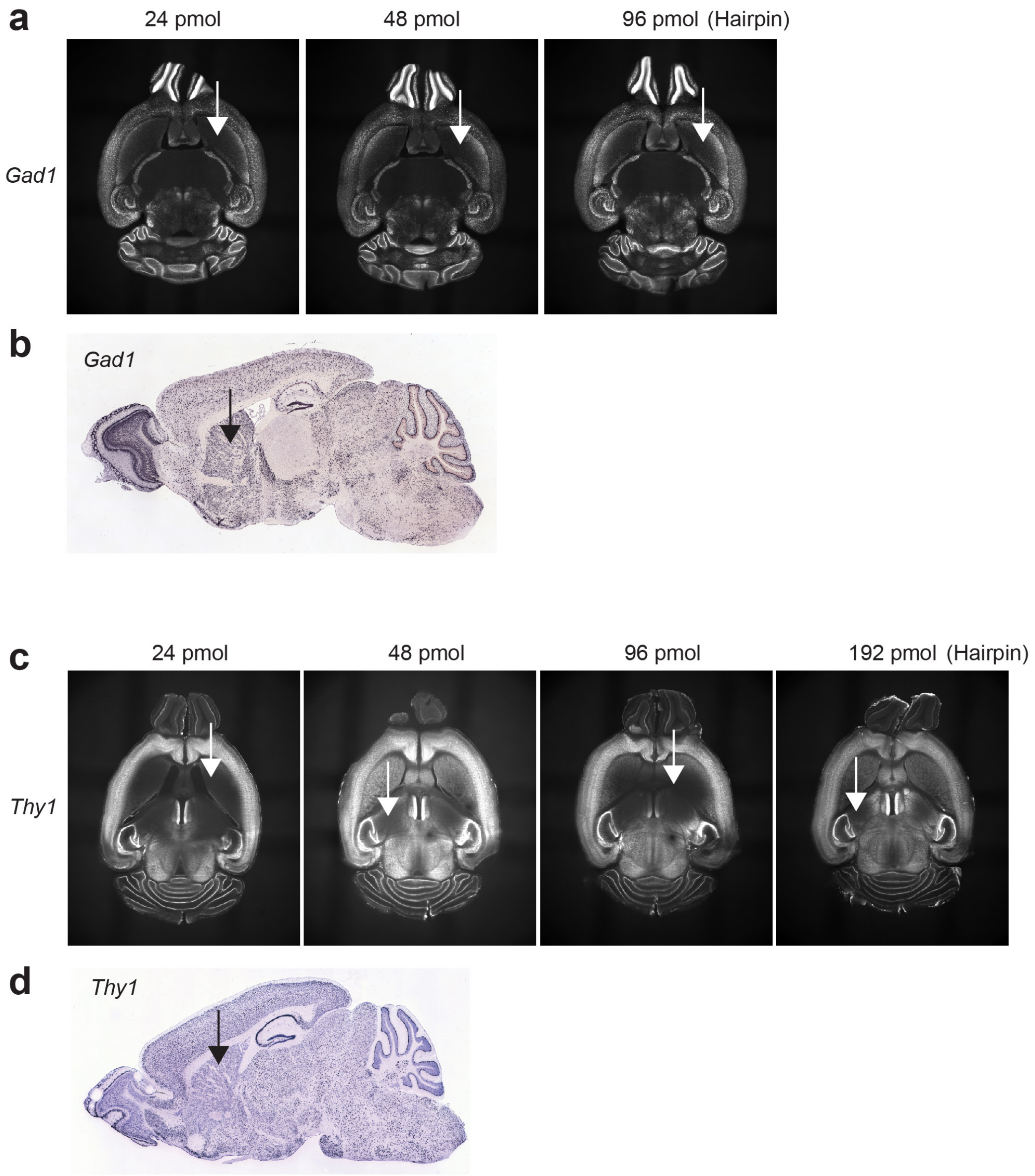

**Supplementary Figure 2 | Assessment of hairpin amount and penetration.** **a**, Representative images of isHCR for *Gad1* using 24 pmol, 48 pmol, and 96 pmol of hairpin DNA. Striatum region showed low penetration and signal intensity in all conditions (arrows). **b**, Allen Brain Atlas image mouse brain section *in situ* hybridized for *Gad1*. Image credit: Allen Institute. Arrow indicates strong staining in striatum. **c**, Representative images of isHCR for *Thy1* using 24 pmol, 48 pmol, 96 pmol, and 192 pmol of hairpin DNA. Random areas of low penetration were observed (arrows). **d**, Allen Brain Atlas image mouse brain section *in situ* hybridized for *Thy1*. Image credit: Allen Institute. Arrow indicates strong staining in striatum.

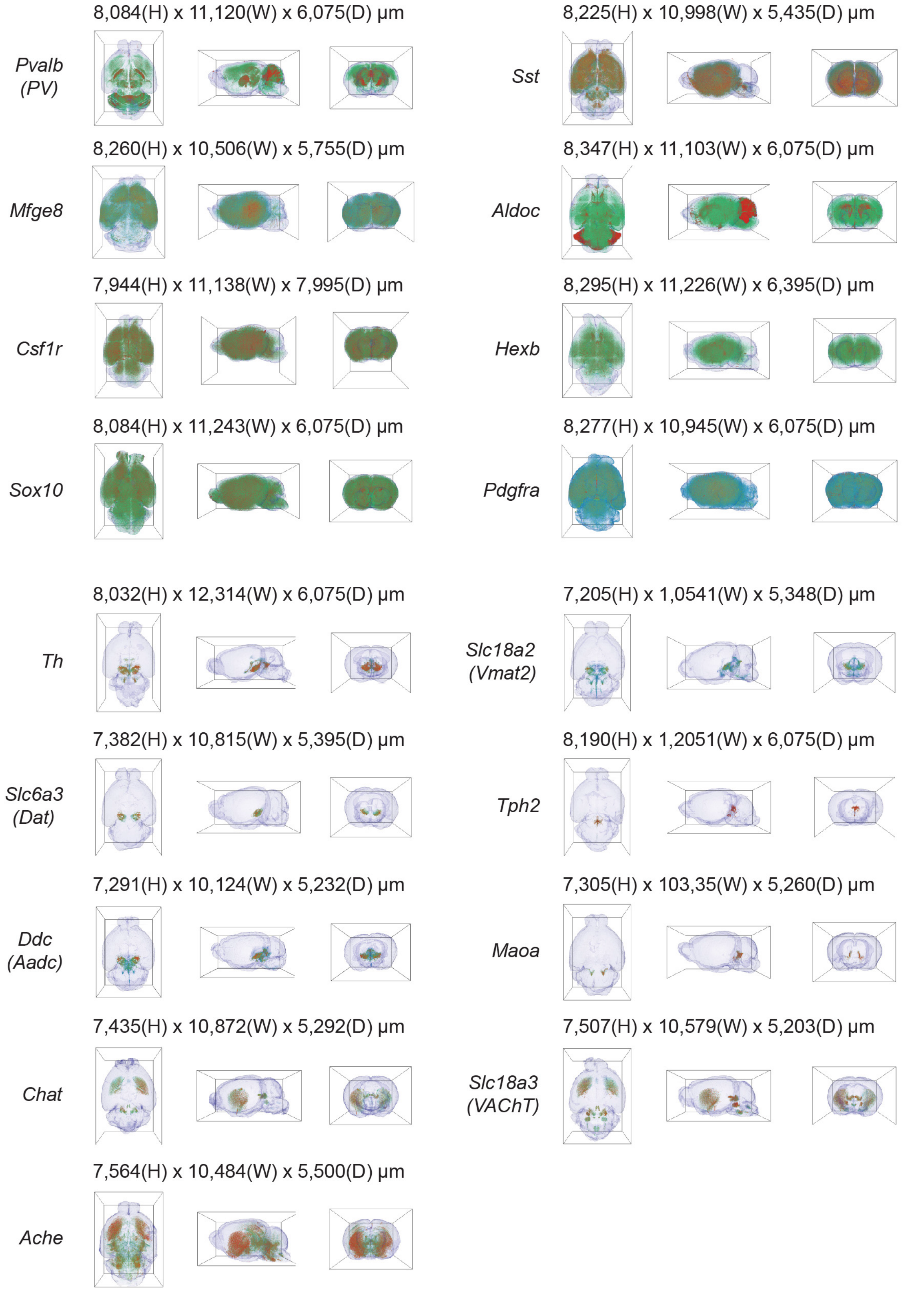

**Supplementary Figure 3 | Volume renderings of whole-brain TRIC-DISCO recordings of various transcripts.** Volume renderings of whole-brain TRIC-DISCO recordings of *Pvalb* (*PV*), *Sst*, *Mfge8*, *Aldoc*, *Csf1r*, *Hexb*, *Sox10*, *Pdgfra*, *Th*, *Slc18a2* (*Vmat2*), *Slc6a3* (*Dat*), *Tph2*, *Ddc* (*Aadc*), *Maoa*, *Chat*, *Slc18a3* (*VAChT*), and *Ache*. The size of each volume rendering (boxed region) is indicated.

#### Supplementary Table Captions

**Supplementary Table 1 | Transcripts used in this study.**
List of custom-designed DNA probes used for whole-brain TRIC-DISCO recordings.
*File: TRIC-DISCO-SuppTable1.xlsx, 70 kB*

#### Supplementary Movie Captions

**Supplementary Movie 1 | Whole-brain multiplex imaging of *Pvalb, Sst*, *Gad1*, and nuclei.**
Z-stack images from a whole-brain TRIC-DISCO multiplex recording of *Pvalb* (green), *Sst* (red), *Gad1* (blue), and nuclei stained with DiYO-1 (gray). The brain was from an eight-week-old mouse.
*File: TRIC-DISCO-SuppMovie1.mp4, 56.3 MB*

**Supplementary Movie 2 | Whole-brain multiplex imaging of *Pvalb, Sst*, *Gad1*, and nuclei in hippocampus.**
Z-stack images from a whole-brain TRIC-DISCO multiplex recording of *Pvalb* (green), *Sst* (red), *Gad1* (blue), and nuclei stained with DiYO-1 (gray). The brain was from an eight-week-old mouse and is the same as in Supplementary Movie 1.
*File: TRIC-DISCO-SuppMovie2.mp4, 31.8 MB*

**Supplementary Movie 3 | Whole-brain imaging of *Pvalb*.**
Z-stack images from a whole-brain TRIC-DISCO recording of *Pvalb*. The brain was from an eight-week-old mouse.
*File: TRIC-DISCO-SuppMovie3.mp4, 57.2 MB*

**Supplementary Movie 4 | Whole-brain imaging of *Sst***
Z-stack images from a whole-brain TRIC-DISCO recording of *Sst*. The brain was from an eight-week-old mouse.
*File: TRIC-DISCO-SuppMovie4.mp4, 53.2 MB*

**Supplementary Movie 5 | Whole-brain imaging of *Mfge8.***
Z-stack images from a whole-brain TRIC-DISCO recording of *Mfge8*. The brain was from an eight-week-old mouse.
*File: TRIC-DISCO-SuppMovie5.mp4, 57.3 MB*

**Supplementary Movie 6 | Whole-brain imaging of *Aldoc.***
Z-stack images from a whole-brain TRIC-DISCO recording of *Aldoc*. The brain was from an eight-week-old mouse.
*File: TRIC-DISCO-SuppMovie6.mp4, 57.0 MB*

**Supplementary Movie 7 | Whole-brain imaging of *Csf1r.***
Z-stack images from a whole-brain TRIC-DISCO recording of *Csf1r.* The brain was from an eight-week-old mouse.
*File: TRIC-DISCO-SuppMovie7.mp4, 45.3 MB*

**Supplementary Movie 8 | Whole-brain imaging of *Hexb.***
Z-stack images from a whole-brain TRIC-DISCO recording of *Hexb.* The brain was from an eight-week-old mouse.
*File: TRIC-DISCO-SuppMovie8.mp4, 58.9 MB*

**Supplementary Movie 9 | Whole-brain imaging of *Sox10.***
Z-stack images from a whole-brain TRIC-DISCO recording of *Sox10.* The brain was from an eight-week-old mouse.
*File: TRIC-DISCO-SuppMovie9.mp4, 55.2 MB*

**Supplementary Movie 10 | Whole-brain imaging of *Pdgfra.***
Z-stack images from a whole-brain TRIC-DISCO recording of *Pdgfra.* The brain was from an eight-week-old mouse.
*File: TRIC-DISCO-SuppMovie10.mp4, 58.2 MB*

**Supplementary Movie 11 | Whole-brain imaging of *Th.***
Z-stack images from a whole-brain TRIC-DISCO recording of *Th.* The brain was from an eight-week-old mouse.
*File: TRIC-DISCO-SuppMovie11.mp4, 51.2 MB*

**Supplementary Movie 12 | Whole-brain imaging of *Slc18a2.***
Z-stack images from a whole-brain TRIC-DISCO recording of *Slc18a2 (Vmat2).* The brain was from an eight-week-old mouse.
*File: TRIC-DISCO-SuppMovie12.mp4, 69.1 MB*

**Supplementary Movie 13 | Whole-brain imaging of *Slc6a3.***
Z-stack images from a whole-brain TRIC-DISCO recording of *Slc6a3 (Dat).* The brain was from an eight-week-old mouse.
*File: TRIC-DISCO-SuppMovie13.mp4, 50.6 MB*

**Supplementary Movie 14 | Whole-brain imaging of *Tph2.***
Z-stack images from a whole-brain TRIC-DISCO recording of *Tph2.* The brain was from an eight-week-old mouse.
*File: TRIC-DISCO-SuppMovie14.mp4, 40.7 MB*

**Supplementary Movie 15 | Whole-brain imaging of *Ddc.***
Z-stack images from a whole-brain TRIC-DISCO recording of *Ddc (Aadc).* The brain was from an eight-week-old mouse.
*File: TRIC-DISCO-SuppMovie15.mp4, 39.5 MB*

**Supplementary Movie 16 | Whole-brain imaging of *Maoa.***
Z-stack images from a whole-brain TRIC-DISCO recording of *Maoa.* The brain was from an eight-week-old mouse.
*File: TRIC-DISCO-SuppMovie16.mp4, 61.3 MB*

**Supplementary Movie 17 | Whole-brain imaging of *Chat.***
Z-stack images from a whole-brain TRIC-DISCO recording of *Chat.* The brain was from an eight-week-old mouse.
*File: TRIC-DISCO-SuppMovie17.mp4, 67.4 MB*

**Supplementary Movie 18 | Whole-brain imaging of *Slc18a3.***
Z-stack images from a whole-brain TRIC-DISCO recording of *Slc18a3 (VAChT).* The brain was from an eight-week-old mouse.
*File: TRIC-DISCO-SuppMovie18.mp4, 67.8 MB*

**Supplementary Movie 19 | Whole-brain imaging of *Ache.***
Z-stack images from a whole-brain TRIC-DISCO recording of *Ache.* The brain was from an eight-week-old mouse.
*File: TRIC-DISCO-SuppMovie19.mp4, 75.8 MB*

**Supplementary Movie 20 | Volume rendering of whole-brain imaging of *Slc18a3*, *Maoa*, *Slc6a3*, *Slc18a2*, *Ache*, *Chat*, *Tph2*, *Ddc*, and *Th*.**
Volume rendering of whole-brain TRIC-DISCO recordings of *Slc18a3* (*VAChT*), *Maoa*, *Slc6a3* (*Dat*), *Slc18a2* (*Vmat2*), *Ache*, *Chat*, *Tph2*, *Ddc* (*Aadc*), and *Th*. The brains were from eight-week-old mice.
*File: TRIC-DISCO-SuppMovie20.mp4, 28.6 MB*

**Supplementary Movie 21 | Volume rendering of whole-brain imaging of *Pvalb.***
Volume rendering of whole-brain TRIC-DISCO recordings of *Pvalb*. The brain was from an eight-week-old mouse.
*File: TRIC-DISCO-SuppMovie21.mp4, 33.8 MB*

**Supplementary Movie 22 | Volume rendering of whole-brain imaging of *Th.***
Volume rendering of whole-brain TRIC-DISCO recordings of *Th*. The brain was from an eight-week-old mouse.
*File: TRIC-DISCO-SuppMovie22.mp4, 26.7 MB*

### Supplementary Methods

**Supplementary Protocol | TRIC-DISCO protocol.**
The whole-brain TRIC-DISCO protocol describing: Reagents, Buffers, Sample Collection, Delipidation and bleaching, Hybridization, Amplification, TRIC, Clearing, and Sample Mounting and Imaging.
*File: TRIC-DISCO-SuppProtocol.pdf, 169 kB*
