## Supplementary Protocol for "Whole-Brain Three-Dimensional Imaging of RNAs at Single-Cell Resolution"

### Whole Brain in Situ HCR
### Supplementary Protocol

**Protocol Contents**


1. **Reagents**
2. **Buffers**
3. **Sample Collection**
4. **Delipidation and bleaching**
5. **Hybridization**
6. **Amplification**
7. **TRIC**
8. **Clearing**
9. **Sample Mounting and Imaging**


---

#### 1. Reagents

| Reagent | Use | Supplier Information |
|---|---|---|
| 30% Hydrogen peroxide solution | Bleaching | Sigma-Aldrich Cat. # 216763 |
| Water | Buffers | MilliQ water, autoclaved |
| Tris(hydroxymethyl) aminomethane | Buffers | VWR Cat. # 71003-490 |
| Formamide (Deionized) | Buffers | Life Technologies Cat. # AM9342 |
| 20x Sodium chloride sodium citrate (SSC) | Buffers | Life Technologies Cat. # 15557-044 |
| Citric acid | Buffers | Merck Millipore Cat. # 100241 |
| Tween 20 | Buffers | Merck Millipore Cat. # 822184 |
| 50x Denhardt's solution | Buffers | Life Technologies Cat. # 750018 |
| Dextran sulfate sodium salt | Buffers | Sigma-Aldrich Cat. # D6001 |
| Heparin | Buffers | Sigma-Aldrich Cat. # H4784 |
| 10X PBS, pH 7.4 | Buffers | Life Technologies Cat. # 70011044 |
| 99.8% Methanol | Dehydration | Sigma-Aldrich Cat. # 179337 |
| Dichloromethane | Delipidation | Sigma-Aldrich Cat. # 270997 |
| Paraformaldehyde | Fixation | Sigma-Aldrich Cat. # 441244 |
| 37% Hydrochloric acid | pH adjustment | Sigma-Aldrich Cat. # 320331 |
| Sodium hydroxide | pH adjustment | Merck Millipore Cat. # 106498 |
| Poly (vinylsulfonic acid, sodium salt) solution, 30% | RNase inhibitor | Sigma-Aldrich Cat # 278424 |
| Dibenzyl ether | RI Matching | Sigma-Aldrich Cat. # 108014 |

### Whole Brain in Situ HCR
### Supplementary Protocol

#### 2. Buffers

**4% Paraformaldehyde (500 mL, store at 4°C or -20 °C)**

| Ingredient | Amount | Final Conc. |
|---|---|---|
| Paraformaldehyde | 20 g | 4 % |
| 10X PBS | 50 mL | 1X |
| 5N Sodium hydroxide | 100 μL | - |
| Water | To 500 mL | - |

*Note:
1) Mix 250ml water and 20g Paraformaldehyde and 100 μL Sodium hydroxide with heating to dissolve Paraformaldehyde.
2) Mix with 10X PBS and 200 mL water and make 500 mL solution in total.

**1M Citric acid, pH 6.0 (100 mL, store at 4°C)**

| Ingredient | Amount | Final Conc. |
|---|---|---|
| Citric acid | 19.2 g | 1 M |
| Sodium hydroxide | Adjust pH to 6.0 | - |
| Water | To 100 mL | - |

**10% Tween 20 (50 mL, kept at RT)**

| Ingredient | Amount | Final Conc. |
|---|---|---|
| Tween 20 | 5 mL | 10% |
| Water | To 50 mL | - |

**50% formamide wash buffer (50 mL, kept at RT)**

| Ingredient | Amount | Final Conc. |
|---|---|---|
| Formamide | 25 mL | 50% |
| 20x SSC | 12.5 mL | 5x |
| 1M citric acid (pH 6.0) | 450 μL | 9 mM |
| 10% Tween 20 | 500 μL | 0.1% |
| 25 mg/mL Heparin | 100 μL | 50 μg/mL |
| 30% PVSA | 500 μL | 1:100 |
| Water | To 50 mL | - |

**50% formamide hybridization buffer (50 mL, kept at 4°C)**

| Ingredient | Amount | Final Conc. |
|---|---|---|
| Formamide | 25 mL | 25% |
| 20x SSC | 12.5 mL | 5x |
| 1M citric acid (pH 6.0) | 450 μL | 9 mM |
| 50x Denhardt's solution | 1 mL | 1x |
| 10% Tween 20 | 500 μL | 0.1% |
| 25 mg/mL Heparin | 100 μL | 50 μg/mL |
| 50% dextran sulfate | 10 mL | 10% |
| 30% PVSA | 500 μL | 1:100 |
| Water | To 50 mL | - |

### Whole Brain in Situ HCR
### Supplementary Protocol

##### Amplification buffer (50 mL, kept at 4°C)

| Ingredient | Amount | Final Conc. |
| --- | --- | --- |
| 20x SSC | 12.5 mL | 5x |
| 10% Tween 20 | 500 µL | 0.1% |
| 50% dextran sulfate | 10 mL | 10% |
| 30% PVSA | 500 µL | 1:100 |
| Water | To 50 mL | - |

##### 5x SSCT (50 mL, kept at RT)

| Ingredient | Amount | Final Conc. |
| --- | --- | --- |
| 20x SSC | 12.5 mL | 5x |
| 10% Tween 20 | 500 µL | 0.1% |
| 30% PVSA | 500 µL | 1:100 |
| Water | To 50 mL | - |

##### 1M Tris-HCl (pH 7.0) (1 L, kept at RT)

| Ingredient | Amount | Final Conc. |
| --- | --- | --- |
| Tris(hydroxymethyl) aminomethane | 25 g | 50% |
| 37% Hydrochloric acid | Adjust pH to 7.0 | |
| Water | To 1 L after pH adjustment | - |

##### 50% dextran sulfate (50 mL, kept at 4°C)

| Ingredient | Amount | Final Conc. |
| --- | --- | --- |
| Dextran sulfate powder | 25 g | 50% |
| Water | To 50 mL | - |

##### 25 mg/mL Heparin (Store at -20°C)

| Ingredient | Amount | Final Conc. |
| --- | --- | --- |
| Heparin | 250 mg | 25 mg/mL |
| Water | To 10 mL | - |

### Whole Brain in Situ HCR
### Supplementary Protocol

#### 3. Sample Collection

i) Collect mouse brain and fix the brain sample in cold 4% PFA overnight at 4°C (with perfusion is preferred).
ii) Dehydrate the sample with series of Methanol from 60% as shown below. Wash duration for whole brain shown here is 1 h. Please adjust accordingly for different samples. Longer incubation, such as 1 day usually does not affect in situ signal.

(Stop point) Here you can store the sample in Methanol at -20°C for long term.

*Note:
1) Follow the local rule of chemical handling for Carcinogenic, mutagenic and reprotoxic chemicals (CMR).
2) Put sample on a shaker with gentle agitation for better buffer exchange.

| | **Duration** | **Buffer** | **Temperature** |
|---|---|---|---|
| **Day 1** | Overnight | 4% Paraformaldehyde | 4°C |
| **Day 2** | 1h | 60% | RT |
| | 1h | 80% | RT |
| | 1h | 100% | RT |
| | Overnight | 100% | RT |

#### 4. Delipidation and bleaching

i) Incubate the sample in 100% DCM for at least one day (Longer incubation does not affect in situ signal).
ii) Wash the sample with 100% Methanol for 1 hour twice at RT.
iii) Incubate the sample in 5% $H_2O_2$ solution in Methanol overnight at 4°C.
iv) Wash the sample with 100% Methanol for 1 hour twice at RT.

(Stop point) Here you can stop and store the sample in Methanol at -20°C for long term.

*Note:
1) Put sample on a shaker with gentle agitation for better buffer exchange.
2) Longer incubation in DCM and Methanol won't affect the signal intensity.

| | **Duration** | **Buffer** | **Temperature** |
|---|---|---|---|
| **Day 3** | Overnight | 100% DCM | RT |
| **Day 4** | 1h | 100% Methanol | RT |
| | 1h | 100% Methanol | RT |
| | Overnight | 5% $H_2O_2$ | 4°C |
| **Day 5** | 1h | 100% Methanol | RT |
| | 1h | 100% Methanol | RT |

#### 5. Hybridization (hybridization oven with rotation is recommended)

i) Incubate the sample in 50% formamide wash buffer with gentle shaking until the sample sinks at RT.
ii) Incubate the sample in 50% formamide wash buffer until the sample sinks at RT.
iii) Block the sample in 50% formamide hybridization buffer at 37°C overnight.
iv) Hybridization in 50% formamide hybridization buffer at 37°C for 3 days.
v) Wash the sample with 50% formamide wash buffer at 37°C for more than 1 hour three times.

*Note:
1) Longer wash time won't usually affect the signal intensity.
2) DiYO-1 can be added in hybridization step in 1:1000 ratio.

| | **Duration** | **Buffer** | **Temperature** |
|---|---|---|---|
| **Day 5** | Until sink | 50% formamide wash buffer | RT |
| | Until sink | 50% formamide wash buffer | RT |
| | Overnight | 50% formamide hybridization buffer | 37°C |
| **Day 6** | 3 days | 50% formamide hybridization buffer | 37°C |
| **Day 9** | 1h | 50% formamide wash buffer | 37°C |
| | 1h | 50% formamide wash buffer | 37°C |
| | 1h | 50% formamide wash buffer | 37°C |

#### 6. Amplification (in situ HCR)

i) Wash the sample with 5XSSCT at RT for more than 1 hour three times.
ii) Equilibrate the sample with amplification buffer overnight at 4°C.
iii) HCR reaction for 3 days at 4°C (400ul solution).
iv) Wash the sample with 5XSSCT at RT for more than 1 hour two times.

*Note: Long incubation of HCR reaction more than 4 days will increase the background noise.

| | **Duration** | **Buffer** | **Temperature** |
|---|---|---|---|
| **Day 9** | 1h | 5XSSCT | RT |
| | 1h | 5XSSCT | RT |
| | 1h | 5XSSCT | RT |
| | Overnight | Amplification buffer | 4°C |
| **Day 10** | 3 days | Hairpin in amplification buffer | 4°C |
| **Day 13** | 1h | 5XSSCT | RT |
| | 1h | 5XSSCT | RT |

### Whole Brain in Situ HCR
### Supplementary Protocol

#### 7. TRIC

i) Wash the sample with 500mM Tris-HCl (pH 7.0) at RT for more than 1 hour three times (last wash is overnight).
ii) Dehydrate the sample with 100% Methanol three times for more than 1 hour at RT (last step is overnight).

(Stop point) Here you can stop and store the sample in Methanol at -20°C in dark place at least for a week.

*Note: Longer wash won't affect the signal intensity.

| | **Duration** | **Buffer** | **Temperature** |
|---|---|---|---|
| **Day 13** | 1h | 500mM Tris-HCl | RT |
| | 1h | 500mM Tris-HCl | RT |
| | Overnight | 500mM Tris-HCl | RT |
| **Day 14** | 1h | 100% Methanol | RT |
| | 1h | 100% Methanol | RT |
| | Overnight | 100% Methanol | RT |

#### 8. Clearing

i) Delipidate the sample in 66%DCM/33%Methanol for 3 hours at RT.
ii) Wash the sample with 100% DCM for 15 minutes twice at RT remove excess Methanol from the tissue.
iii) Transfer the sample to DBE solution and incubate overnight at RT.
iv) (Optional) Change the DBE solution to fresh (then transparency increases a little).

| | **Duration** | **Buffer** | **Temperature** |
|---|---|---|---|
| **Day 15** | 3h | 66%DCM/33%Methanol | RT |
| | 15 min | 100% DCM | RT |
| | 15 min | 100% DCM | RT |
| | Overnight | DBE | RT |

#### 9. Sample Mounting and Imaging

i) Prepare sample holders for mounting if you need.
ii) Mount sample on the holder and move the holder into imaging chamber of LaVision Ultramicroscope II.

*Note: Take image within 2 weeks because the signal intensity gradually decreases.
*Comment: C57BL/6 sample is better fit to LaVision Ultramicroscope II because of the size of the brain.
